# Activation of a zinc metallochaperone by the alarmone ZTP

**DOI:** 10.1101/594382

**Authors:** Pete Chandrangsu, Xiaojuan Huang, John D. Helmann

## Abstract

Bacteria tightly regulate intracellular zinc levels to ensure sufficient zinc to support essential functions, while preventing toxicity. The bacterial response to zinc limitation includes the expression of putative zinc metallochaperones belonging to subfamily 1 of the COG0523 family of G3E GTPases. However, the client proteins and the metabolic processes served by these chaperones are unclear. Here, we demonstrate that the *Bacillus subtilis* YciC zinc metallochaperone (here renamed ZagA for <u>Z</u>TP <u>a</u>ctivated <u>G</u>TPase <u>A</u>) supports *de novo* folate biosynthesis under conditions of zinc limitation through direct interaction with the zinc dependent GTP cyclohydrolase, FolE. Furthermore, we identify a role for the alarmone ZTP, a modified purine biosynthesis intermediate, in the response to zinc limitation. ZTP, a signal of 10-formyl-tetrahydrofolate deficiency (10f-THF) in bacteria, transiently accumulates as the Zn dependent GTP cyclohydrolase FolE begins to fail and stimulates the interaction between ZagA and FolE to sustain folate synthesis despite declining zinc availability.

**Importance:** Metallochaperones provide a mechanism for cells to regulate the delivery of metals to newly synthesized apoproteins. By selectively targeting specific proteins for metallation, cells can ensure that key pathways remain functional even as metals become limiting for growth. The COG0523 family of proteins contain a subgroup of candidate metallochaperones (the YciC subfamily) induced under conditions of zinc limitation. Although YciC family proteins have been suggested to be GTP-dependent metallochaperones, specific interactions with client proteins have not been demonstrated. Here, we show that the *Bacillus subtilis* YciC (renamed ZagA) protein responds to ZTP as an activating ligand rather than GTP, and interacts specifically with a Zn-dependent enzyme critical for folate synthesis (FolE). Thus, under conditions of Zn limitation ZagA is synthesized, and as folate synthesis fails, it selectively delivers Zn to FolE to sustain folate synthesis.

## Introduction

Transition metals are required for life and participate as cofactors in a wide range of essential biological functions. Of these, zinc is often considered a “first among equals” as it serves as cofactor for ~4-10% of all proteins (1). As such, zinc plays a key role in host-microbe interactions (2). In a process termed nutritional immunity, the host may restrict bacterial access to zinc in response to infection through the production of calprotectin, an S100 protein produced by cells of the immune system (3).

The physiological states associated with zinc homeostasis can be generally described as excess, sufficiency, deficiency and limitation (or starvation) (4). Excess zinc can lead to toxic consequences, and leads to the expression of protective mechanisms including sequestration or efflux. Sufficiency refers to the optimal zinc concentration to support zinc dependent cellular processes. Deficiency is characterized by decreased growth, altered metabolism, and deployment of an adaptive response. As zinc levels fall further, zinc limitation results as defined by the failure of essential zinc dependent processes and cessation of growth.

Bacteria utilize complex mechanisms to respond to metal stress. In *Bacillus subtilis*, a model Gram-positive bacterium, zinc homeostasis is maintained by the coordinated action of two DNA binding metalloregulators: Zur, the sensor of zinc sufficiency, and CzrA, the sensor of zinc excess. Under conditions of zinc sufficiency, the dimeric Fur family metalloregulator Zur binds DNA in its zinc-loaded form and represses transcription (5). Genes repressed by Zur are derepressed in three distinct groups as cells transition from sufficiency to limitation (6). This sequential regulation is facilitated, in part, by negative cooperativity between the two zinc sensing sites, one in each subunit of the Zur dimer (7).

During the initial response to zinc limitation, zinc independent paralogs of the L31 and L33 ribosomal proteins (L31* and L33* r-proteins, respectively) are expressed (6, 8, 9). The ribosome is proposed to contain 6-8 equivalents of zinc (10). Given that cells may contain >30,000 copies of the ribosome during rapid growth, the ribosome represents a substantial zinc storage pool. Two of these zinc containing r-proteins, L31 and L33, are loosely associated with the surface of the ribosome and are non-essential for translation (11-13). Expression of the Zur-regulated L31* and L33* r-proteins, which do not require zinc for function, facilitates displacement of their zinc-associated paralogs (L31 and L33) thereby enabling mobilization of ribosome-associated zinc. The expression of alternative ribosomal proteins under zinc limitation is a conserved feature in a variety of bacteria (14, 15), and provides a fitness advantage when zinc is limited (11, 16, 17). This mobilization response precedes the expression of high affinity uptake systems in both *B. subtilis* (6) and *Salmonella* Typhimurium (18).

If cells experience continued zinc starvation, cells shift their adaptive response from zinc mobilization to zinc acquisition. During this phase, cells derepress the genes encoding the ZnuABC high affinity uptake system and the YciC protein, a putative zinc metallochaperone (here renamed ZagA for <u>Z</u>TP <u>a</u>ctivated <u>G</u>TPase <u>A</u>) (6). ZagA is a member of the zinc-associated subfamily 1 of the COG0523 family G3E GTPases (19). COG0523 proteins are evolutionarily related to well characterized nickel metallochaperones, including UreG for urease and HypB for nickel hydrogenase (20). The functions of COG0523 proteins, which are found in all domains of life, are generally associated with the assembly or function of metalloproteins. COG0523 family metallochaperones have been identified with functions related to cobalt (CobW), iron (Nha3) and zinc (YeiR and ZigA) homeostasis (19). However, the functions of COG0523 GTPases with respect to zinc homeostasis are poorly understood. GTPase and zinc-binding activities have been reported for both *Escherichia coli* YeiR and *Acinetobacter baumannii* ZigA(21, 22). ZigA is postulated to help activate a zinc-dependent histidine ammonia-lyase, HutH, which is implicated in the mobilization of a histidine-associated zinc pool (22).

As zinc levels are depleted further and essential zinc dependent processes begin to fail, genes encoding zinc-independent functions are derepressed to compensate and allow for survival. In *B. subtilis*, derepression of a zinc-independent S14 paralog (RpsNB) ensures continued ribosome synthesis if the zinc-containing paralog can no longer access zinc required for proper folding and function (12). S14 is an early assembling r-protein and is essential for de novo ribosome synthesis. Furthermore, derepression of a zinc independent GTP cyclohydrolase (FolEB) supports continued folate biosynthesis under conditions where the constitutively expressed, but zinc dependent FolE enzyme fails (23). The order of the adaptive responses to declining zinc levels in *B. subtilis* is mobilization (from ribosomal proteins), acquisition, and finally replacement of zinc-dependent functions (e.g. S14) with non-zinc containing paralogs (6). This same order of response is also predicted from an analysis of Zur-binding affinities in *Salmonella* Typhimurium (18).

Here, we demonstrate that zinc limitation results in failure of the folate biosynthetic pathway due to a loss of FolE activity, and this results in a transient purine auxotrophy that can be partially overcome by the eventual derepression of FolEB. At the onset of zinc limitation, the purine biosynthetic intermediate 5-aminoimidazole-4-carboxamide ribonucleotide (AICAR), also known as ZMP, accumulates and is phosphorylated to generate ZTP. The Zur-regulated metallochaperone ZagA is activated by ZTP to deliver zinc to FolE to sustain folate synthesis. These results suggest that a subset of zinc-associated COG0523 proteins are activated by ZTP, rather than GTP, and provide an example of a physiologically relevant ZTP-receptor protein.

## Results

### Zinc deficient cells experience folate starvation

The physiological consequences of zinc starvation are unclear, and given the ubiquity of zinc as a cofactor for protein-folding and catalysis the precise physiological processes that fail are not immediately obvious. The strongest hints come from a close examination of the Zur regulon in diverse bacteria, which often include zinc-independent paralogs of zinc-dependent proteins (14-16). These zinc-independent proteins are generally thought to maintain cellular functions that are normally carried out by zinc-dependent proteins, which may fail under conditions of zinc limitation.

In *B. subtilis*, the Zur-dependent regulation of a zinc-independent GTP cyclohydrolase (FolEB) suggests that folate biosynthesis may represent a major metabolic bottleneck caused by zinc limitation, at least in cells lacking an alternate enzyme. To determine if folate biosynthesis is also compromised during zinc limitation of wild-type cells, we compared sensitivity to EDTA, a potent metal chelator known to result in zinc limitation in *B. subtilis* when grown in minimal medium (5). During growth in minimal medium, the folate biosynthetic pathway is active and 50 *μ*M EDTA elicited growth inhibition which could be reversed by addition of inosine **(Fig 1A)**. This suggests a failure of purine biosynthesis, which is known to be a major consequence of folate limitation. Moreover, cells lacking *folE2*, and therefore completely reliant on FolE for *de novo* folate biosynthesis, were significantly more sensitive to EDTA inhibition than wild-type **(Fig 1B)**. These results suggest that zinc limitation results in folate deficiency due to failure of the zinc dependent enzyme FolE, and this can limit growth even when the alternative, zinc-independent FolE (FolEB) can be induced to compensate for FolE failure.

**Figure 1.**
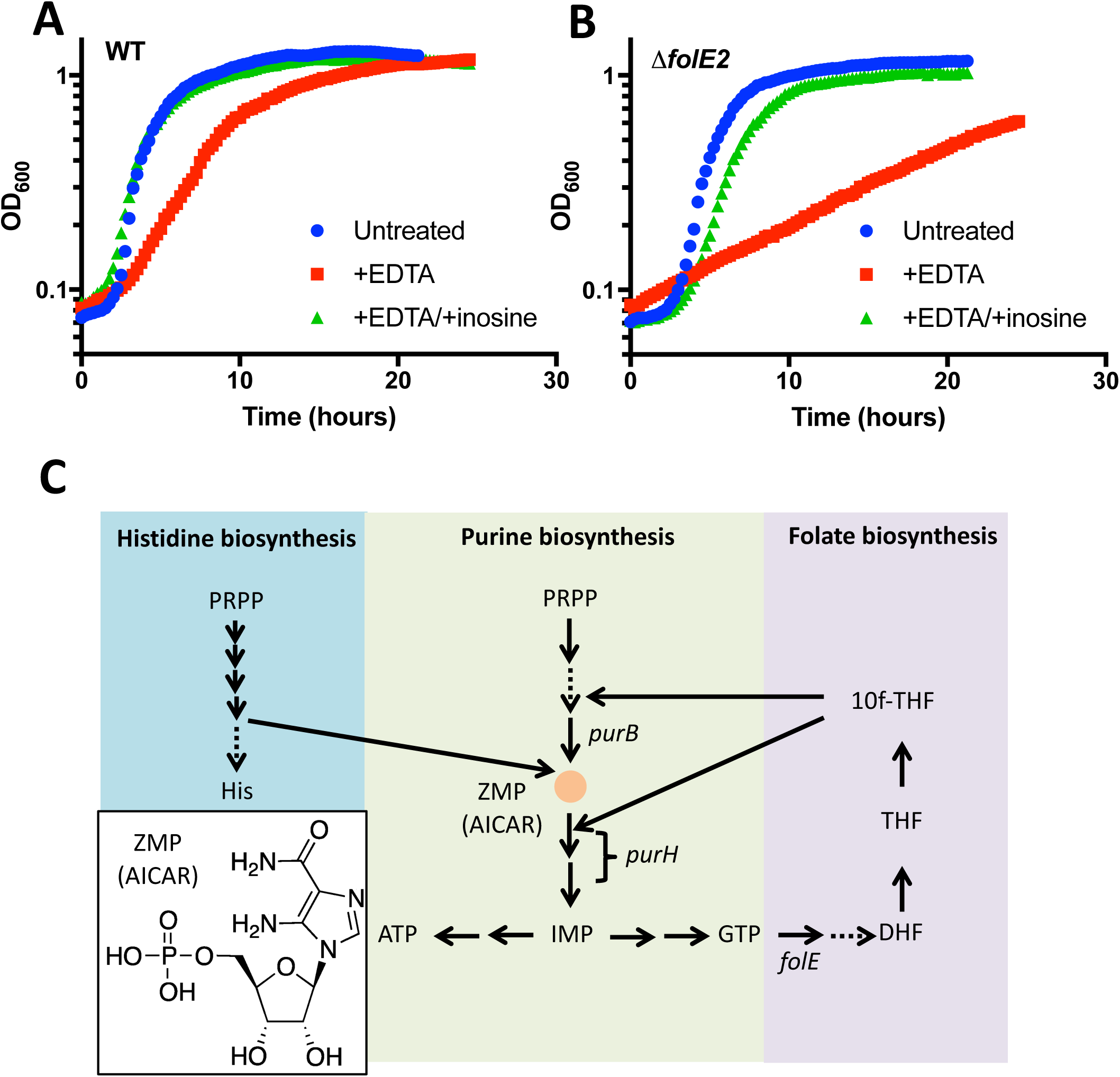
Purine biosynthesis is the major metabolic bottleneck caused by folate limitation during zinc starvation. Growth curves of wild type (A) and a *folE2* mutant (B) in the presence or absence of EDTA (50 mM) or inosine (100 mM) supplementation. (C) Diagram of ZMP producing pathways. Dashed arrows indicate multiple steps. Abbreviations: PRPP=phosphoribosyl pyrophosphate, His=histidine, DHF=dihydrofolic acid, THF=tetrahydrofolate, 10f-THF=10-formyl tetrahydrofolate, AICAR=5-Aminoimidazole-4-carboxamide ribonucleotide.

Folate derived cofactors, such as tetrahydrofolate (THF), are required for a number of cellular processes. Inhibition of THF biosynthesis leads to cell death as a result of purine auxotrophy and consequent thymine deficiency (“thymineless death”) (24). In *B. subtilis*, purine biosynthesis is the primary bottleneck caused by 10f-THF depletion after treatment with antifolates, such as trimethoprim (TMP) (25). 10f-THF is used as a formyl group donor at two steps in purine biosynthesis **(Fig 1C).** As also noted in prior studies, 10f-THF deficiency leads to the failure of the later step required for inosine monophosphate (IMP) production, the common precursor to ATP and GTP (26). This critical step is catalyzed by PurH, a bifunctional enzyme that utilizes 10-formyl-tetrahydrofolate as a formyl donor to convert AICAR (aminoimidazole carboxamide ribonucleotide or ZMP) into IMP.

### Accumulation of Z nucleotides (ZMP/ZTP) protects cells from zinc starvation

In the course of these studies, we unexpectedly observed that a *purH* mutant is more resistant than wild-type to zinc limitation when grown on rich medium (LB) **(Fig. 2A)**. The phenotypes associated with disruption of *purH* may result from a general inability to produce purines and/or the accumulation of the IMP precursor, ZMP. To distinguish between these models, we generated a strain lacking *purB*, which is immediately upstream of *purH* in the purine biosynthetic pathway and catalyzes ZMP production **(Fig 1C)**. We reasoned that if the contribution of *purH* to zinc homeostasis requires ZMP, the phenotypes associated with loss of *purH* would be abrogated in the absence of *purB*. Indeed, a *purB* mutant is more sensitive to EDTA than wild-type, and this effect is dramatically enhanced in a strain also lacking the ZnuABC zinc uptake system **(Fig 2A,B)**. These data link ZMP to zinc homeostasis and suggest that accumulation of ZMP, or the resultant ZTP, may protect cells against zinc limitation.

**Figure 2.**
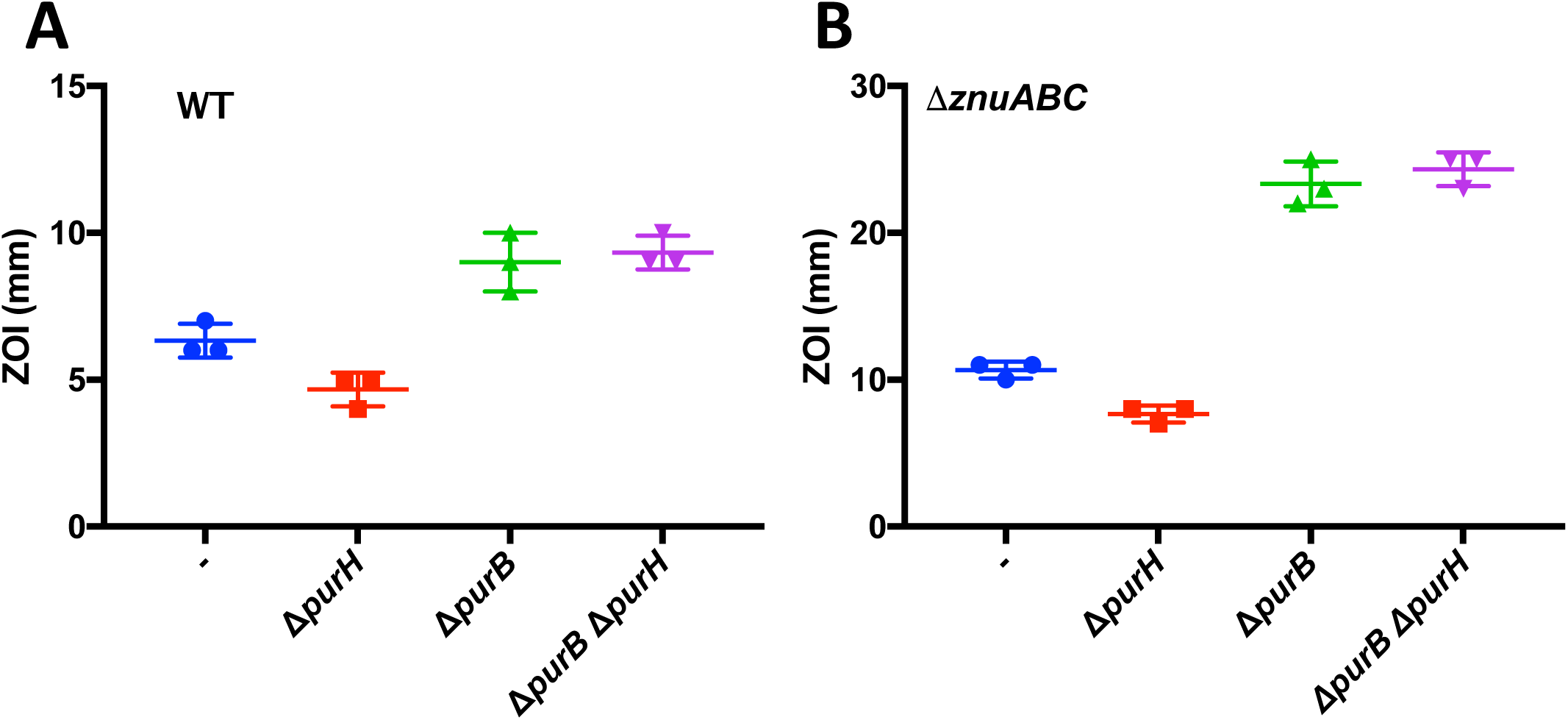
ZMP accumulation protects cells from zinc starvation. EDTA sensitivity of *purH*, *purB* and *purB purH* mutants in wild-type (A) or *znuABC* mutant (B) backgrounds as measured by disk diffusion assay.

### ZTP serves as a signal of folate deficiency during zinc limitation

Disruption of *purH* or loss of 10f-THF production is predicted to result in the accumulation of the purine intermediate ZMP (or AICAR) **(Fig 1C)**. ZMP is phosphorylated to produce ZTP, an “alarmone” proposed to act as a signal of 10f-THF deficiency by Bochner and Ames in 1982 (26). The functional consequence of ZTP accumulation remained a mystery for over thirty years until the discovery of ZTP sensing riboswitches that regulate expression of genes which ensure sufficient 10f-THF to support purine biosynthesis (27). Our data suggest that ZMP/ZTP is also linked to the *B. subtilis* response to zinc limitation, despite the lack of any known ZMP/ZTP-sensing riboswitches in the *B. subtilis* 168 strain. We hypothesized that zinc limitation may induce folate deficiency thereby leading to an accumulation of ZMP/ZTP, which then mediates an increased resistance to zinc depletion by an unknown mechanism.

To evaluate intracellular Z nucleotide levels during zinc depletion, we monitored expression of a *lacZ* reporter construct under the control of the *B. subtilis SG-1 pfl* riboswitch **(Fig 3A**) (27). Since the *pfl* riboswitch does not distinguish between ZMP and ZTP, this reporter provides an estimate of total Z nucleotide levels (27). Z nucleotide binding to the riboswitch aptamer domain prevents the formation of a transcription termination stem-loop structure located upstream of the translation start site **(Fig 3A)**. We speculated that zinc deprivation induced by EDTA would result in an increase in reporter expression. Indeed, we observed induction of the *pfl-lacZ* reporter in the presence of an EDTA impregnated disk **(Fig 3B)**.

**Figure 3.**
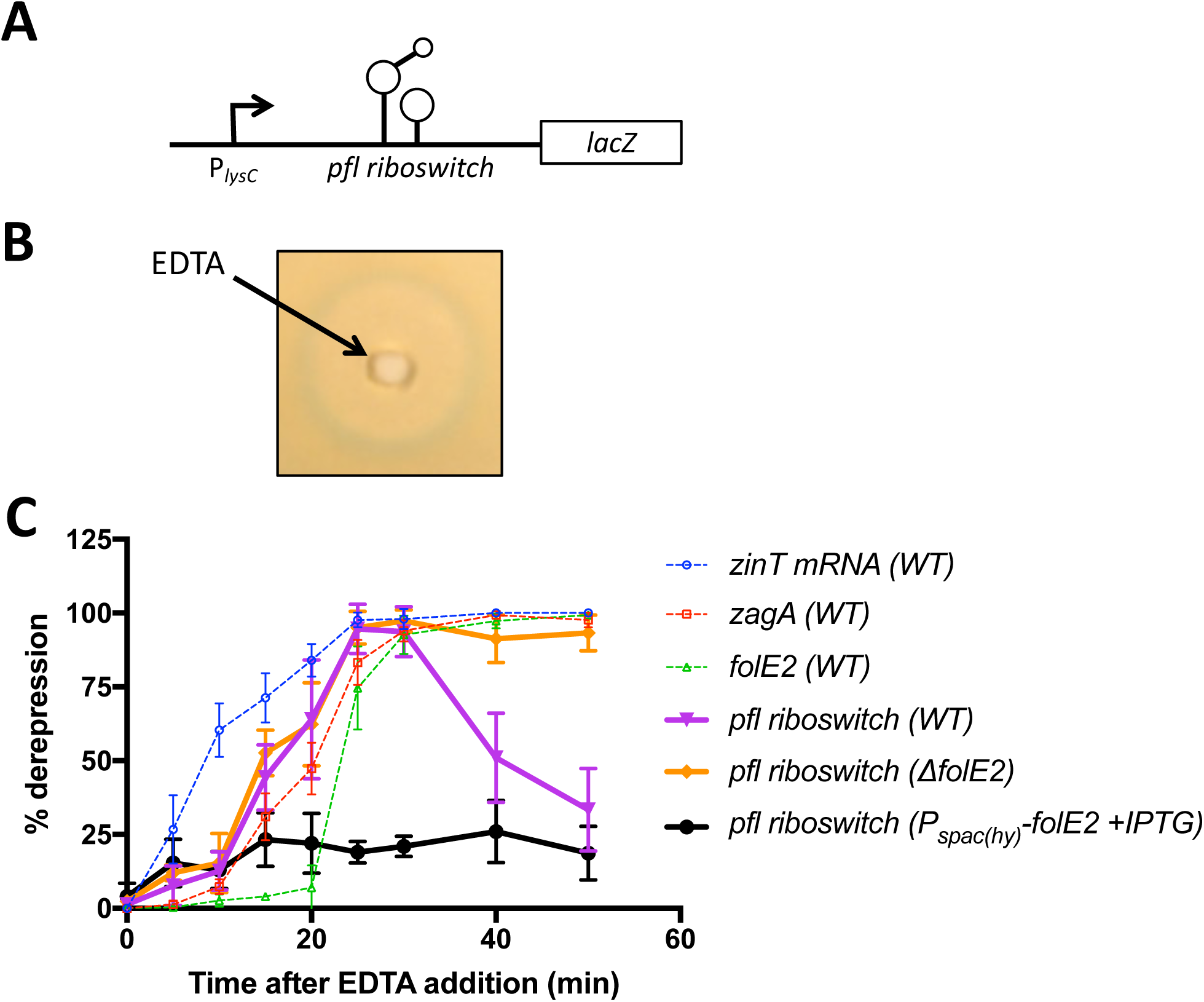
Z nucleotides accumulate under conditions of zinc starvation. (A) Schematic representation of the *pfl* riboswitch-*lacZ* reporter construct. (B) Induction of the *pfl* riboswitch-*lacZ* fusion in response to an EDTA impregnated disk. (C) Induction of the *pfl* riboswitch-*lacZ* fusion and derepression of the Zur regulon as a function of time after EDTA (250 mM) addition.

As monitored by the activity of the *pfl-lacZ* reporter, ZMP/ZTP accumulation by EDTA was transient, reaching a maximum after ~20 minutes of exposure to 250 μM EDTA **(Fig 3C)**. We note that the induction of the *pfl* riboswitch commenced after induction an early induced Zur-repressed gene (*zinT*), and correlated in time with the induction of the middle gene, *zagA* (formerly *yciC*), as monitored by RT-PCR. Interestingly, the subsequent decrease in *pfl-lacZ* reporter expression was correlated with the derepression of the late gene, *folE2* **(Fig 3C)**. We reasoned that the restoration of *de novo* folate biosynthesis by FolE2 would restore PurH activity and thereby consume ZMP. Together with turnover of ZTP, this would lead to a loss in activation of the *pfl* riboswitch. Indeed, expression from the *pfl* riboswitch remained elevated in the absence of *folE2* **(Fig 3C)**. Additionally, constitutive expression of *folE2* prior to zinc limitation prevented the accumulation of Z nucleotides **(Fig 3C)**. These data are consistent with our hypothesis that zinc limitation results in a failure of folate biosynthesis due to a loss of PurH activity and Z nucleotide accumulation.

### *B. subtilis* ZagA protects cells from zinc starvation and requires Z nucleotides

The consequences of Z nucleotide accumulation on zinc homeostasis are not well understood. One possibility is that Z nucleotides may directly interact with zinc and serve as an intracellular zinc buffer. However, ZTP does not bind zinc with high affinity **(Fig S1)**. Alternatively, Z nucleotides may serve as a signal for zinc limitation. The only known Z nucleotide receptor is the recently described *pfl* riboswitch (27), which is not present in the laboratory strain of *B. subtilis* 168 used in our studies. This motivated us to consider alternative possibilities for Z nucleotide effectors.

Given the close association of the *pfl* riboswitch with folate biosynthetic genes in other bacteria (27), we surmised that ZTP accumulation (e.g. in a *purH* mutant) might facilitate growth under zinc limiting conditions by affecting folate biosynthesis. In many bacteria, *folE2* is located in close chromosomal association with a COG0523 protein, a Zur-regulated GTPase proposed to deliver zinc to proteins under conditions of zinc limitation (19). We therefore speculated that YciC, a *B. subtilis* COG0523 protein, might function as a <u>Z</u>TP-<u>a</u>ssociated <u>G</u>TPase (ZagA) to deliver zinc to one or more client proteins. Consistent with this hypothesis, a *zagA* mutant is more sensitive to EDTA than wild type **(Fig 4A)**. Moreover, the EDTA resistance of a *purH* mutant, which accumulates Z nucleotides, is abrogated when *zagA* is deleted **(Fig 4A)**. Additionally, the effect of mutation of *zagA* and *purB* nucleotides, is not additive **(Fig 4A)**. This indicates that the increased resistance to zinc deprivation in the *purH* strain, which accumulates ZMP/ZTP, requires ZagA. Finally, we note that in a strain (*purB*) unable to make Z nucleotides, *zagA* no longer has a discernable role in resistance to zinc deprivation. Similar results were seen in a *znuABC* mutant background **(Fig S2)**. The greater sensitivity of the *purB* mutant relative to the *zagA* may suggest that Z nucleotides also have ZagA-independent roles.

**Figure 4.**
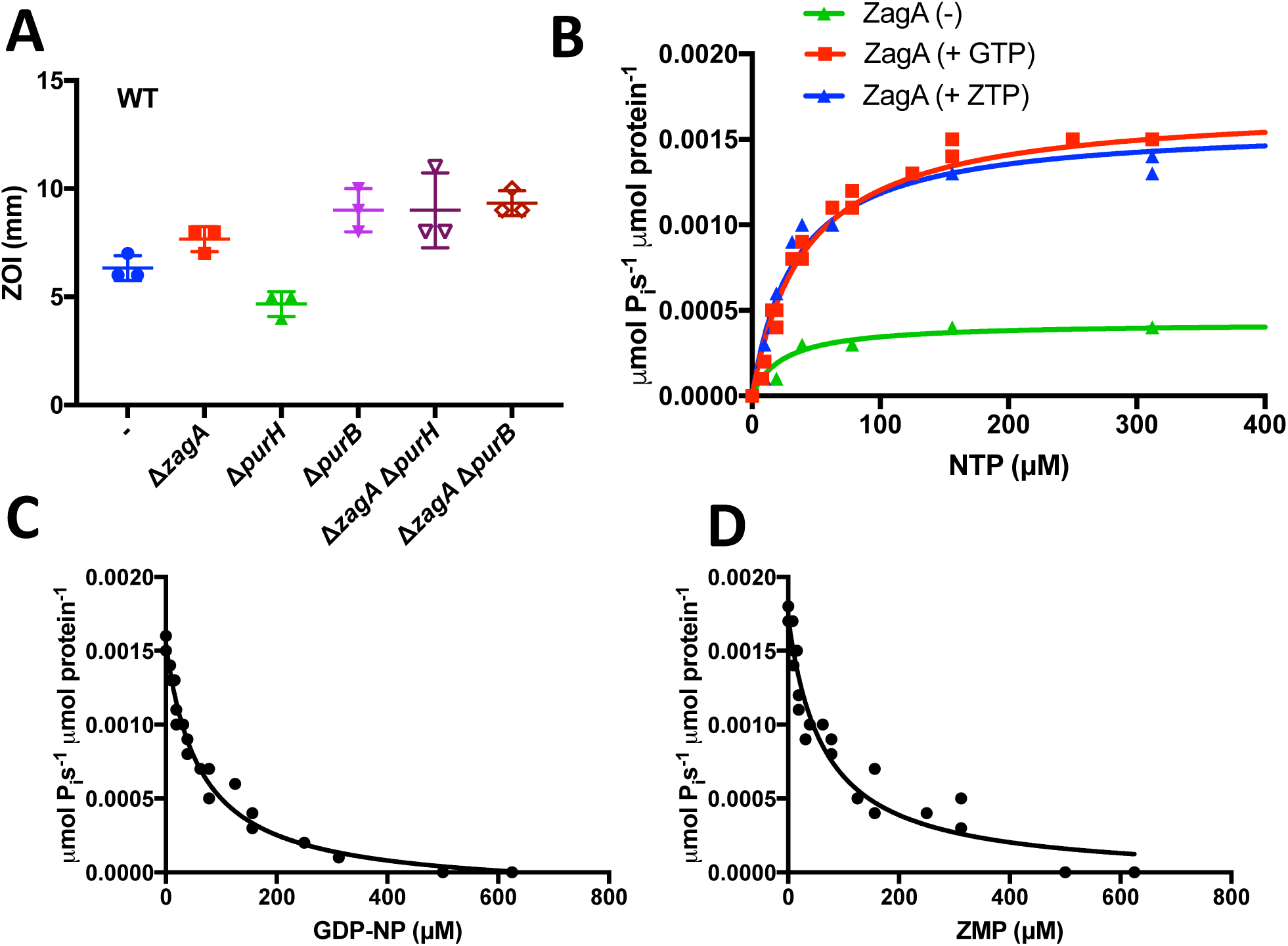
ZagA hydrolyzes ZTP. (A) EDTA sensitivity of *zagA*, *purH*, *purB*, *zagA purH*, and *zagA purB* mutants as measured by disk diffusion assay. (B) ZagA nucleotide hydrolysis activity as measured by the Malachite green assay. (C) Inhibition of ZagA ZTPase activity by addition of the non-hydrolyzable GTP analog, GDP-NP. (D) Inhibition of ZagA GTPase activity by addition of ZMP.

### ZagA is a both a GTPase and a ZTPase

ZagA is a member of the COG0523 family of G3E GTPases, which have been shown to hydrolyze GTP *in vitro* (21, 22). Indeed, under our conditions, ZagA is a GTPase with an apparent K_m_ for GTP of 40 μM, consistent with that measured for other COG0523 proteins **(Fig 4B)**. Since intracellular ZMP and ZTP levels rise to levels at or near GTP levels (~4x for ZMP; ~0.8x for ZTP) upon folate starvation (26), and in light of the structural similarity between ZTP and GTP, we hypothesized that ZagA may also interact with and hydrolyze Z nucleotides. Indeed, ZagA is a ZTPase with an apparent K_m_ for ZTP (36 μM) comparable to that of GTP **(Fig 4B)**. Furthermore, GDP-NP, a non-hydrolyzable analog of GTP, inhibited ZTP hydrolysis **(Fig 4C)**. Conversely, ZMP inhibited GTP hydrolysis **(Fig 4D)**. These data indicate that both ZTP and GTP are ZagA substrates.

### Z nucleotides trigger ZagA interaction with FolE to sustain folate synthesis during zinc limitation

Information regarding client proteins served by COG0523 family proteins is limited. Given the synteny and zinc-dependent coregulation of *zagA* and *folE2*, we postulated that ZagA might physically interact with either FolEB or the zinc-dependent FolE proteins. However, in initial studies using a bacterial two-hybrid assay, no interaction was observed with either protein. Since ZagA can hydrolyze ZTP *in vitro*, we reasoned that the putative ZagA interaction with its client proteins may require Z nucleotides. Therefore, we reassessed the interaction in cells grown on minimal medium where the purine biosynthetic pathway is active and Z nucleotides are produced. In addition, we utilized the folate biosynthesis inhibitor trimethoprim (TMP) to induce Z nucleotide accumulation. Interaction between ZagA and FolE was only detected when the cells were treated with the antifolate TMP **(Fig 5A, B)** and the strength of this interaction increases in a concentration dependent manner **(Fig S3)**. In contrast, no interaction between ZagA and FolE2 was observed **(Fig 5A)**. These data support a model where ZagA functions as a chaperone to deliver zinc to FolE in a ZTP dependent manner. By sustaining FolE activity, ZagA and ZTP serve to delay the failure of folate biosynthesis under conditions of declining zinc availability.

**Figure 5.**
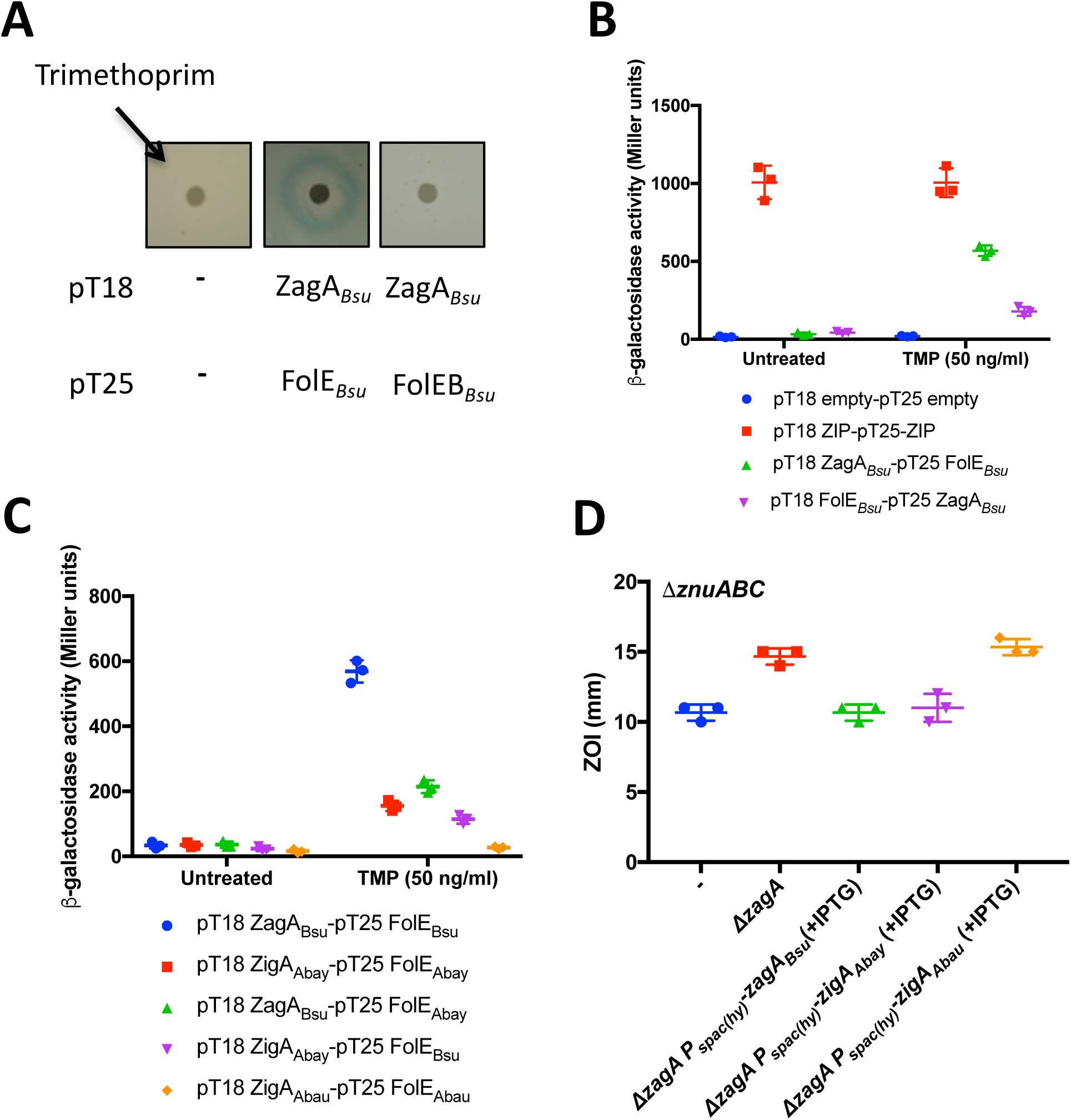
Z nucleotide accumulation stimulates the YciC-FolE interaction. (A) Disk diffusion assays of *E. coli* strains containing the ZagA, FolE, FolE2 bacterial two hybrid constructs in the presence of trimethoprim (TMP). (B) *β*-galactosidase activity of the ZagA and FolE bacterial two hybrid constructs after 30 min of treatment with TMP. (C) *β*-galactosidase activity of the ZagA and FolE from *B. subtilis* (*Bsu*)*, A. baumanii* (*Abau*), or *A. baylyi* (*Abay*) bacterial two hybrid constructs after 30 min of treatment with TMP. (D) Complementation of the EDTA sensitivity of *B. subtilis zagA* mutant with *B. subtilis* or *A. baylyi zigA* or *A. baumanii zigA*.

Since Zur regulated COG0523 family proteins are often encoded in close proximity with the gene encoding FolEB, we reasoned that the interaction of COG0523 family proteins and FolE may be broadly conserved. Using this bacterial two hybrid assay, we detected significant interaction between the *Acinetobacter baylyi* ZagA homolog and its FolE in the presence of TMP **(Fig 5C)**. Furthermore, *A. baylyi* ZagA interacts with *B. subtilis* FolE, and *B. subtilis* ZagA with *A. baylyi* FolE **(Fig 5C)**.

*Acinetobacter baumanii* encodes a distinct COG0523 family member, ZigA, that is postulated to function in the metallation of histidine ammonium lyase (22). No significant interaction was observed between ZigA and FolE2 of *Acinetobacter baumanii*, nor can *A. baumanii* ZigA interact with *A. baylyi* or *B. subtilis* FolE **(Fig 5C)**. Additionally, ectopic expression of *A. baylyi* ZagA, but not *A. baumanii* ZigA, is able to complement the EDTA sensitivity of a *B. subtilis zagA* mutant **(Fig 5D)**. These data suggest that ZagA-related COG0523 proteins sustain FolE-dependent folate biosynthesis under zinc deficiency (leading to *zagA* induction) and when folate synthesis fails (as signaled by Z nucleotide accumulation). Moreover, this adaptive response is likely present in many bacteria, and other COG0523 proteins, such as ZigA (22), likely have related functions, but with different client proteins.

To directly assess the impact of ZagA on FolE function, we monitored FolE GTP cyclohydrolase I activity as a function of time after exposure to a subinhibitory concentrations of EDTA (250 μM) to induce zinc deficiency **(Fig 6)**. A fluorescence based assay in which the conversion of GTP to dihydroneopterin triphosphate was used to monitor FolE activity in crude cell extracts (28). After 30 minutes of exposure to EDTA, FolE activity decreased slightly in wild type and near full activity was recovered after 60 minutes, presumably to Zur-regulon derepression and FolE2 expression. Consistent with this hypothesis, restoration of FolE activity was not observed in cell extracts prepared from strains lacking *folE2*. In strains lacking ZagA, FolE activity decreases dramatically compared to wild-type after 30 min before recovering, which suggests that ZagA supports FolE activity under conditions of zinc deficiency.

**Figure 6.**
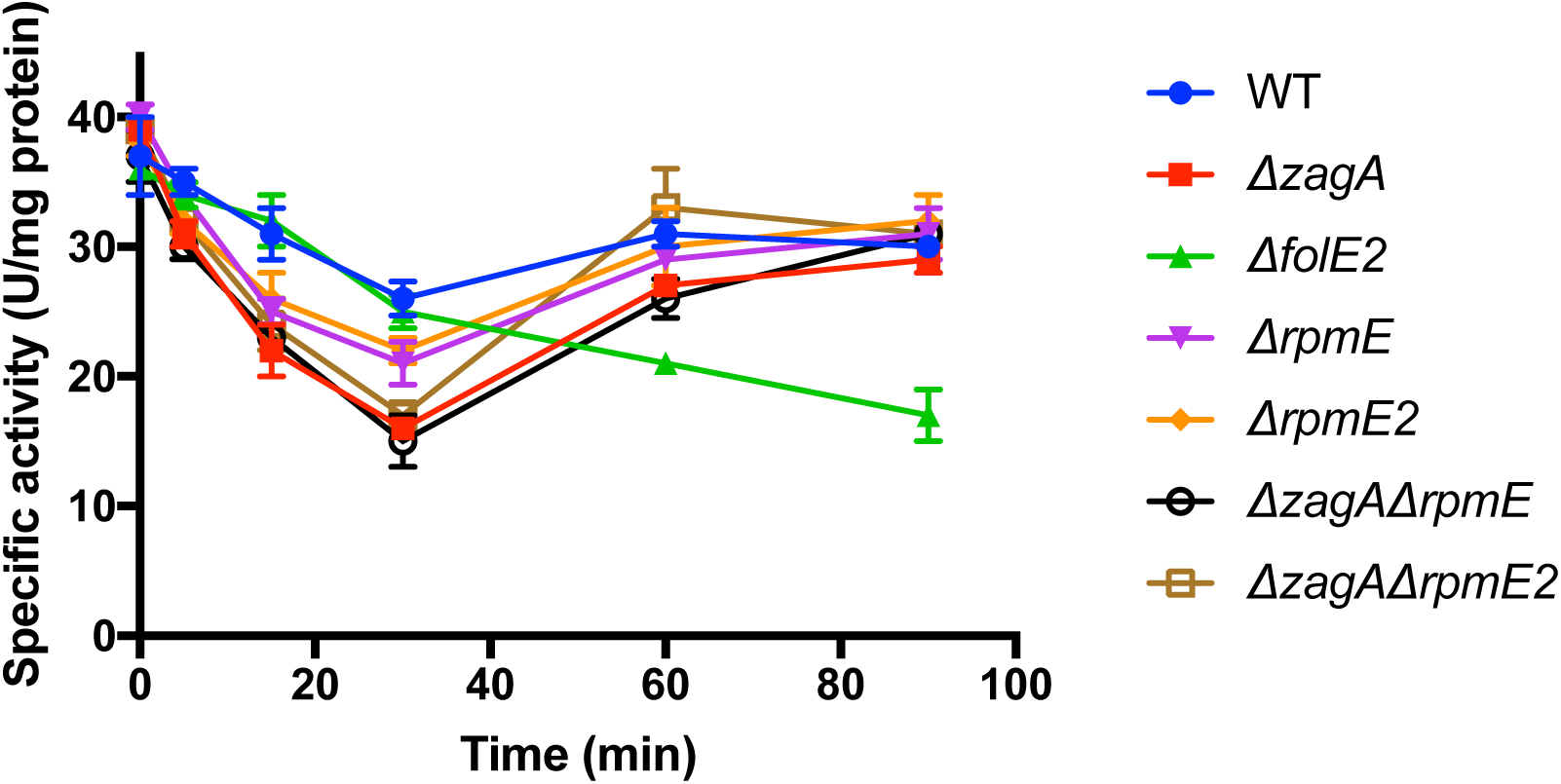
ZagA can access a ribosome associated zinc pool to support FolE GTP cyclohydrolase activity. GTP cyclohydrolase I activity in crude cell lysates of *B. subtilis* WT, *zagA*, *folE2*, *rpmE*, *rpmE2*, *zagA rpmE*, and *zagA rpmE2* mutants as measured by the conversion of GTP to neopterin triphosphate as monitored by fluorescence (265 nm excitation, 450 emission).

### ZagA accesses a ribosome-associated zinc pool to support FolE function

The initial response of *B. subtilis* to zinc limitation is the derepression of two alternative ribosomal proteins (L31* and L33*) that can displace their zinc-containing paralogs from the surface of the ribosome (6, 13). The release of L31 and L33 is postulated to mobilize a pool of bioavailable zinc to sustain critical zinc-dependent enzymes. We therefore set out to quantify the contribution of L31 and L33 to the cellular zinc quota and to test whether ZagA relies on this pool of mobilizable zinc to sustain FolE function.

To quantify the mobilizable zinc pool associated with ribosomal proteins, we measured total intracellular zinc and the zinc content of purified ribosomes in several genetic backgrounds. Our results indicate that ribosome-associated zinc (0.19 mM) accounts for ~20% of total cellular zinc (0.88 mM), with ~5.6 ± 0.9 zinc ions per ribosome (n=6) when cells are grown in rich LB medium. In strains missing the zinc containing L31 (*rpmE*) and L33 ribosomal proteins (encoded by *rpmGA* and *rpmGB*), total cellular zinc and ribosomally associated zinc is reduced to 0.73 mM and 0.12 mM respectively. As expected, the estimated zinc per ribosome in strains lacking L31 and L33 decreases by ~2 (~3.6 ± 0.8 zinc ions per ribosome; n=4). Under these growth conditions, we estimate a total content of ribosomes of 2.5±0.5 × 10^4^ per cell. Thus, the mobilization of zinc from the ribosome can potentially redistribute ~5 x 10^4^ zinc atoms per cell (depending on total ribosome content per cell at the onset of zinc deficiency), which represents a substantial pool of zinc to sustain growth.

To monitor the impact of ribosome-associated zinc on the intracellular bioavailable zinc pools we took advantage of the ability of Zur to serve as a bioreporter. The *folEB* gene is only induced when zinc levels fall to growth limiting levels as one of the last genes induced during zinc depletion (6). We therefore fused the Zur-regulated *folEB* promoter to an operon encoding luciferase and monitored gene expression in response to zinc depletion elicited with EDTA **(Fig S4)**. In wild-type cells we failed to observe induction from the *folEB* promoter, even with concentrations of EDTA (5 *μ*M and 10 *μ*M) that slowed growth. In contrast, cells lacking the gene encoding L31* (*ytiA*, also renamed as *rpmEB*), displayed a strong induction from the *folEB* promoter, despite displaying an overall similar response to EDTA in terms of growth inhibition. This suggests that induction of L31* is required to mobilize zinc from the ribosome and that, in so doing, it delays the decrease in cellular zinc levels that is required for derepression of the *folEB* promoter. Cells lacking the zinc-containing L31 protein (*rpmE*) were much more sensitive to growth inhibition by EDTA **(Fig S4)** and displayed a very strong transcriptional induction of *folEB* even at the lowest tested levels of EDTA. These results suggest that cells lacking L31 are much more easily depleted of zinc, and this effect is stronger than in cells lacking L31*. One interpretation of this result is that L31* stimulates the mobilization of zinc from L31, but may not be absolutely required for cells to access this zinc pool. We note that in *B. subtilis* 168 strains, the corresponding zinc mobilization system involving the L33 proteins is often inactive due to a frame-shift mutation in the gene (*rpmGC*) encoding the zinc-independent paralog (L33*). This likely contributes to the strong phenotypes noted here due to disruption of the L31*/L31 zinc mobilization response.

We hypothesized that zinc mobilized from the ribosome may be utilized by ZagA to support FolE activity under conditions of zinc limitation. We therefore monitored the decline in FolE GTP cyclohydrolase activity in extracts from strains lacking either the L31 (*rpmE*) or L31*(*rpmE2*) proteins after treatment with EDTA **(Fig 6)**. In both cases, FolE activity declined more rapidly within the first 20 minutes of EDTA treatment when compared to wild-type. Additionally, the effect of *zagA* and *rpmE* or *rpmE2* were not additive, which suggests that ZagA and the ribosomal proteins function in the same pathway. These data support a model in which zinc mobilized from the surface of the ribosome by the earliest induced proteins (including L31* and when present L33*) can then be used by ZagA to support FolE function, and thereby delay the eventual induction of the alternative enzyme FolEB.

## Discussion

Accumulation of Z nucleotides as a result of folate limitation has been linked to diverse metabolic consequences. In mammals, ZMP (or AICAR) is able to inhibit the proliferation of many types of cancer cells due to the activation of AMP-activated protein kinase, a regulator of the cellular response to metabolic imbalances (29). In bacteria, ZMP is known to be an allosteric inhibitor of enzymes involved in gluconeogenesis (fructose-1,6-bisphosphatase) and coenzyme A biosynthesis (pantoate β-alanine ligase) (30, 31). However, the impact of ZTP, the triphosphorylated ZMP derivative, on cellular physiology is less well understood.

Over 30 years ago, ZTP was proposed to act as a signal of 10f-THF deficiency (26). Only recently, with the recent discovery of the ZTP sensing *pfl* riboswitch, was ZTP accumulation shown to influence purine and folate biosynthesis gene expression (27). To date, no protein target for ZTP has been identified. Here, we describe a role for the ZTP alarmone in activation of the ZagA zinc metallochaperone. ZagA is a ZTPase, and we suggest that ZTP is likely required for delivery of zinc to FolE, as supported by our bacterial two-hybrid studies, and perhaps to other client proteins.

The role of Z nucleotides is coordinated with the transcriptional response (regulated by Zur in *B. subtilis*) to zinc limitation **(Fig 7)**. When *B. subtilis* experiences zinc deficiency, folate biosynthesis begins to fail due to a decrease in the activity of the zinc dependent GTP cyclohydrolase FolE and this elicits the accumulation of ZMP/ZTP. Concurrently, the ZagA metallochaperone is derepressed which can respond to ZTP by binding FolE, presumably for zinc delivery, thereby allowing for continued FolE activity and a restoration of folate biosynthesis. Eventually, as cells transitions from zinc deficiency to limitation, expression of the zinc independent FolE isozyme, *folE2*, is derepressed. FolEB allows for continued folate biosynthesis even as FolE fails.

**Figure 7.**
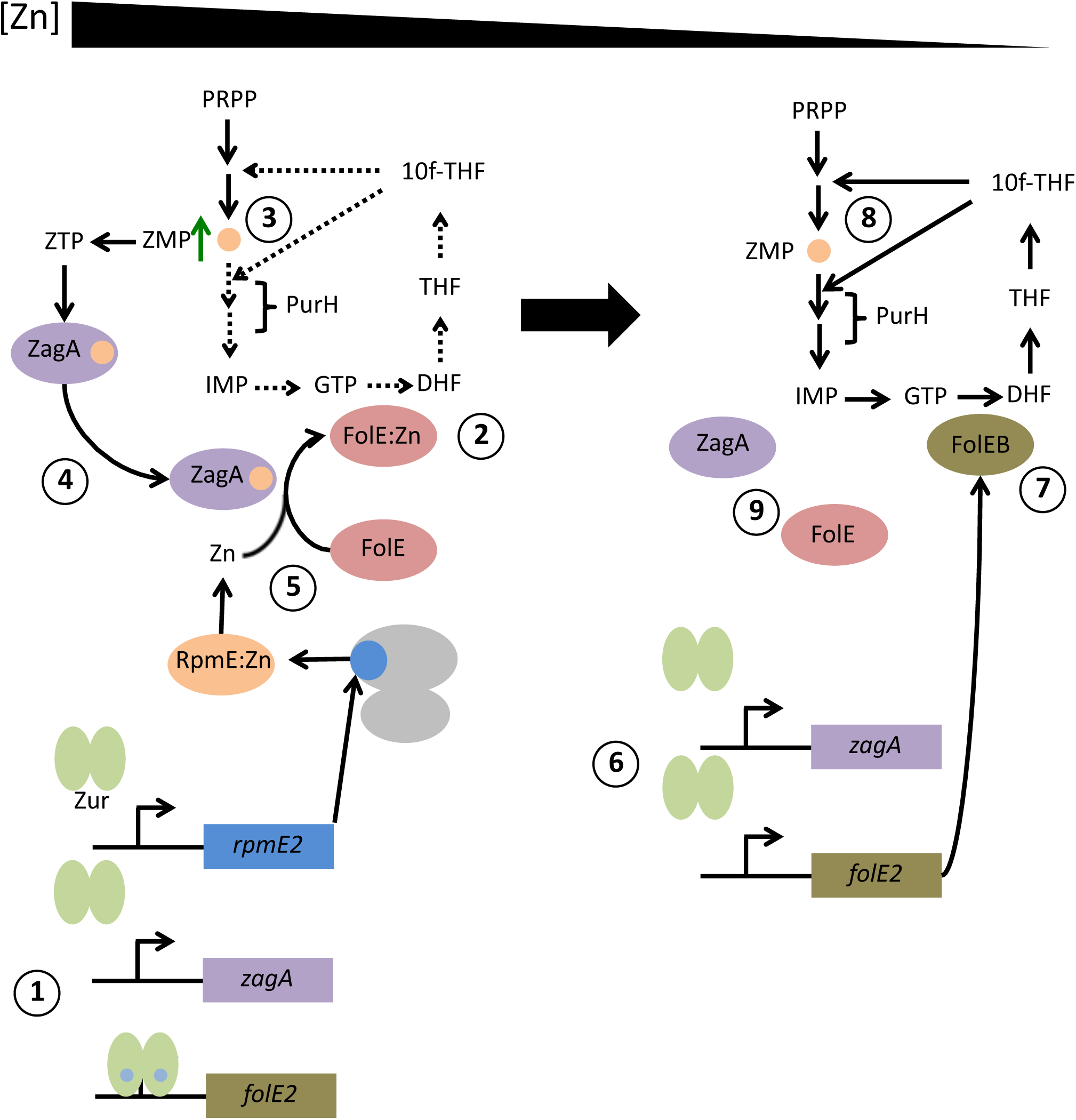
Proposed model of the role of Z nucleotides in the response to zinc limitation. As cells experience zinc limitation, the Zur regulon is derepressed in three distinct waves. The first set of genes to be derepressed (omitted for clarity) includes the zinc independent r-protein paralog L31* (*rpmEB*). (1) L31* can then displace the zinc containing L31 r-protein from the ribosome. As zinc availability continues to decrease, (2) *zagA* (formerly *yciC*) expression is induced. Concurrently, (3) FolE activity begins to decline leading to a decrease in 10f-THF, the substrate for the purine biosynthetic enzyme PurH. As a result, (4) ZMP accumulates and is converted to ZTP. (5) ZTP stimulates ZagA activity and allows for ZagA interaction with FolE, which allows for continued folate production in the presence of zinc limitation. (6) If cells, continue to experience zinc limitation, the final set of Zur regulated genes is derepressed, which includes *folEB*, encoding a zinc independent paralog of FolE. (7) FolEB is able to functionally replace the inactive FolE and, as a result, (8) ZMP levels decline as the purine biosynthetic pathway is functional.

Metallochaperones play a central role in metal homeostasis by delivering metal cofactors to their cognate proteins, thereby providing metal specificity as well as preventing toxicity associated with free cytosolic metal ions (20). The ZagA zinc metallochaperone belongs to subfamily 1 of the COG0523 family of G3E GTPases, proteins associated with the maturation of metal dependent proteins. COG0523 proteins are related to well characterized metallochaperones for nickel, including UreG (for urease) and HypB (for hydrogenase). The first characterized COG0523 protein characterized was *Pseudomonas denitrificans* CobW, which is proposed to contribute to the delivery of cobalt into the cobalamin (Vitamin B12) cofactor (32). A second class of COG0523 proteins is comprised of nitrile hydratase (NHase) activators that facilitate the hydration of nitriles to amides utilizing either iron or cobalt (33).

The third class of COG0523 proteins is related to zinc homeostasis as hinted by their regulation by the zinc sensing metalloregulator, Zur. ZigA, a Zur regulated COG0523 protein from *A. baumanni*, is suggested to deliver zinc to histidine lyase thereby modulating cellular histidine levels, an intercellular zinc buffer (22). Recent results suggest that *A. baumanni zigA* mutants grown in conditions of zinc and iron depletion, as imposed by calprotectin, experience flavin rather than folate limitation. Flavin synthesis in this organism can be initiated by RibA, a Zn-dependent GTP cyclohydrolase II (GCHII), which appears to fail under conditions of Zn limitation (34). However, whether ZigA helps to metallate RibA and/or other specific client proteins is not yet established.

Our data suggest that the *B. subtilis* COG0523 protein, ZagA, is able to hydrolyse ZTP, as well as GTP **(Fig 4B)**. Under folate limiting conditions, cellular ZTP and GTP levels are nearly equal, while ZMP levels accumulate dramatically. Thus, *in vivo*, ZagA and related enzymes may function with either nucleotide. We speculate that as Z nucleotide levels accumulate under zinc limiting conditions, and prior to the derepression of *folE2*, ZagA may utilize Z nucleotides preferentially. As folate biosynthesis is restored upon *folE2* derepression and Z nucleotide levels decrease **(Fig 3B)**, ZagA may continue to function with GTP rather than ZTP. It is even possible, although highly speculative, that ZagA could recognize different client proteins depending on the bound nucleotide.

Genomic analysis offers insight into the cellular processes where ZagA and related metallochaperones may be required. ZagA-like proteins are often encoded near or within operons containing paralogs of zinc dependent proteins (19). Interestingly, the ZagA client protein is not the neighboring FolEB paralog, but rather the zinc containing FolE protein **(Fig 5A)**. In other organisms, proteins predicted to fail under zinc starvation include those involved in heme, pyrimidine and amino acid biosynthesis. For instance, *Pseudomonas aeruginosa* encodes DksA2, a zinc independent paralog of DksA, which is an RNAP binding transcription factor required for appropriate response to amino acid starvation (the “stringent” response) (16). DksA contains a structural zinc binding site, whereas DksA2 does not. Thus, DksA2 can functionally substitute for DksA under conditions of zinc limitation or thiol stress (35, 36). By analogy with our observation that ZagA interacts with FolE, it is reasonable to hypothesize that the *P. aeruginosa* COG0523 protein may interact with the zinc containing DksA to ensure that the cell can mount an effective stringent response. Additionally, the link between DksA and COG0523 proteins also suggests a possible role for the alarmone, guanosine tetraphosphate (ppGpp) in the response to zinc limitation. Thus, the processes that fail as zinc levels become limiting for growth will likely be organism dependent and the proper delivery of zinc to the most critical client proteins may be determined by both the expression of specific COG0523 GTPases and their ability to respond to cellular effectors such as ZTP and perhaps other nucleotide alarmones.

## Materials and Methods

### Strains and growth conditions

Strains used in this study are listed in Table S1. Bacteria were grown in the media described in the following sections. When necessary, antibiotics were used at the following concentrations: chloramphenicol (3 μg ml^-1^), kanamycin (15 μg ml^-1^), spectinomycin (100 μg ml^-1^), and tetracycline (5 μg ml^-1^). Additionally, metal starvation was induced by the addition of EDTA at the concentrations indicated. Markerless in-frame deletion mutants were constructed from BKE strains as described previously (Koo et al., 2013). Briefly, BKE strains were acquired from the Bacillus Genetic Stock Center, chromosomal DNA was extracted, and the mutation, containing an *erm* cassette, was transformed into our wild-type (WT) strain 168. The *erm* cassette was subsequently removed by the introduction of plasmid pDR244, which was later cured by growing at the nonpermissive temperature of 42°C. Gene deletions were also constructed using long flanking homology PCR and chromosomal DNA transformation was performed as described (37).

### Gene expression analysis

Cells were grown at 37°C in MOPS-based minimal medium medium supplemented with 1% glucose and 20 amino acids (50 μg ml^-1^) with rigorous shaking till OD_600_ ~0.4. 1 ml aliquots were treated with 1 mM EDTA for the indicated amount of time. Total RNA from both treated and untreated samples were extracted RNeasy Mini Kit following the manufacturer’s instructions (Qiagen Sciences, Germantown, MD). RNA samples were then treated with Turbo-DNA free DNase (Ambion) and precipitated with ethanol overnight. RNA samples were re-dissolved in RNase-free water and quantified by NanoDrop spectrophotometer. 2 μg total RNA from each sample was used for cDNA synthesis with TaqMan reverse transcription reagents (Applied Biosystems). qPCR was then carried out using iQ SYBR green supermix in an Applied Biosystems 7300 Real Time PCR System. 23S rRNA was used as an internal control and fold-changes between treated and untreated samples were plotted.

### EDTA sensitivity assays

For disk diffusion assays, strains were grown in LB at 37°C with vigorous shaking to an OD_600_~0.4. A 100 μl aliquot of these cultures was added to 4 ml of LB soft agar (0.7% agar) and poured on to prewarmed LB agar plates. The plates were then allowed to solidify for 10 minutes at room temperature in a laminar flow hood. Filter disks (6 mm) were placed on top of the agar and 5 *μ*l of EDTA (500 mM) was added to the disks and allowed to absorb for 10 minutes. The plates were then incubated at 37°C for 16-18 hours. The diameter of the zone of inhibition was measured. The data shown represent the values (diameter of the zone of inhibition minus diameter of the filter disk) and standard deviation of three biological replicates.

### Bacterial two hybrid assay

The bacterial two-hybrid assay was performed as described previously (38). ZagA, FolE, FolE2 from *B. subtilis, A. baumanii* or *A. baylyi* and ZigA from *A. baumanii* were fused to the T18 or T25 catalytic domains of adenylate cyclase. Co-transformed strains of *E. coli BTH101* expressing combinations of T18 and T25 vectors were plated on LB agar and incubated at 30°C for 48 hours. One milliliter of LB medium, supplemented with 100 μg ml^−1^ of ampicillin, 50 μg ml^−1^ of chloramphenicol and 0.5 mM of IPTG, was inoculated and incubated at 30°C to an OD_600_~0.4. One hundred microliters of the culture was mixed with prewarmed 4 ml of M9 medium supplemented with 1% glucose, 10 μg ml^-1^thiamine, appropriate antibiotics, 0.5 mM IPTG and 40 μg ml^−1^ Xgal. containing 0.75% agar. The soft agar was poured onto prewarmed M9 medium plates (1.5% agar) supplemented with 1% glucose, 10 μg/ml thiamine containing appropriate antibiotics, 0.5 mM IPTG and 40 μg ml^−1^ Xgal. A Whatman filter disk impregnated with 5 μM of 50 mg ml^-1^ of trimethoprim was placed on the agar. The plates were incubated at 30°C overnight.

For quantitative β-galactosidase assays, cells were grown in M9 medium supplemented with 1% glucose, 10 μg ml^-1^ thiamine, appropriate antibiotics, 0.5 mM IPTG at 30°C from OD_600_ ~0.02 to OD_600_ ~0.4. One ml of culture was removed to tubes on ice containing 4 ml of Z buffer (0.06 M Na_2_HPO_4_.7 H_2_O, 0.04 M NaH_2_PO_4_.H_2_O, 0.01 M KCl, 0.001 M MgSO_4_, 0.05 M β–mercaptoethanol) for at least 10 min and lysed by sonication. β-galactosidase activity was determined as described previously.

### Overexpression and purification of ZagA

The *zagA* (*yciC*) gene was cloned using primers YciC-LIC-F: TACTTCCAATCCAATGCTATGAAAAAAATTCCGGTTACCGT and YciC-LIC-R: TTATCCACTTCCAATGCTATTGATTCAGCTTCCATTTAA and cloned in pMCSG19c using ligation independent cloning according to (39). The resulting clone was transformed into *E. coli* BL21(DE3) pLysS (40). One liter of liquid LB with 200 μg ml^-1^ ampicillin was inoculated with 1 ml of overnight culture and grown at 37°C to OD_600_ of 0.4. The culture was cooled down to room temperature, IPTG was added to 0.3 mM, and then the culture was incubated at 14°C with shaking for 9 hours. Cells were collected by centrifugation and stored at −80° C. ZagA was purified using Ni-NTA beads (Prepease Histidine purification beads, Life Technologies) according to the manufacturer’s recommendations. ZagA protein was further purified using an FPLC Superdex 200 sizing column using the buffer system, 50 mM Tris-HCl pH 8.0, 150 mM NaCl and 10% glycerol and stored at −80°C.

### GTPase activity assay

GTPase activity was measured by the Malachite green assay (Sigma). Briefly, purified ZagA (1 μM) was incubated with 0–1 mM GTP in assay buffer A in a volume of 90 μL. After 90 min, 35 μL of buffer B was added, incubated for 3 min, and reaction stopped by addition of 15 μL 35% citric acid (Sigma) in 4 N HCl. After 30 min, the absorbance at 680 nm was measured and the concentration of free phosphate was calculated using a standard curve.

### GTP cyclohydrolase activity assay

GTP cyclohydrolase I activity was assessed in crude cell extracts essentially as previously described (28). This assay measures the formation of dihydroneopterin triphosphate from GTP. Crude cell extracts were incubated in a buffer containing 100 mM Tris-HCl pH8.5, 2.5 mM EDTA pH 8.0, 1 mM DTT, and 1 mM GTP (0.5 ml total reaction volume) at 42°C for 30 minutes. At the end of the reaction, an equal volume of activated charcoal (40 μg ml^-1^) was added. The mixture was filtered through a 0.22 μM syringe filter and washed sequentially with 5 ml of water, 5 ml of 5% ethanol, and 5 ml of 50% ethanol/3.1% NaOH. The concentration of neopterin triphosphate in the final wash was determined by fluorescence (265 nm excitation, 450 nm emission).

### Preparation of crude ribosomes

*Bacillus subtilis* crude ribosomes were purified as previously described (23). Briefly, 500 ml of an OD_600_ of ~0.4 LB or MM cultures were harvested and resuspended in buffer I (10 mM Tris [pH 7.6], 10 mM magnesium acetate, 100 mM ammonium acetate, 6 mM β-mercaptoethanol (BME), 2 mM phenylmethylsulfonyl fluoride [PMSF]). Cells were then disrupted by a French press, after removal of cell debris, the supernatant was centrifuged for at 45,000 rpm and 4°C for 100 min in a Thermal Scientific Sorvall MTX 150 micro-ultracentrifuge. The precipitate was dissolved in buffer II (20 mM Tris [pH 7.6], 15 mM magnesium acetate, 1 M ammonium acetate, 6 mM BME, 2 mM PMSF) and centrifuged at 18,000 rpm for 60 min at 4°C in a Thermal Scientific Sorvall MTX150 micro-ultracentrifuge. Then 2 ml aliquots of supernatant were layered onto 2 ml of buffer II containing a 30% (w/v) sucrose bed and centrifuged at 45,000 rpm for 3.5 h at 4°C. The precipitate was resuspended in buffer III (50 mM Tris-HCl, pH 8.0, 6 mM β-mercaptoethanol and 2 mM PMSF). Concentrations of purified ribosomes were quantified by absorbance (1 *A*260 = a 26 nM concentration of 70S ribosomes), and protein composition of the purified crude ribosomes are analyzed by mass spectrometry. Copies of ribosome per cell were calculated by combining ribosome concentrations, cell numbers and culture volume. Measurements were made with six independent preparations for wild-type (CU1065) and four preparations for the CU1065 derivative lacking the Zn-containing L31 and L33 proteins (HB19657). Note that *B. subtilis* 168 contains two genes (*rpmGA* and *rpmGB*) encoding Zn-containing L33 proteins, and one pseudogene for a Zn-independent homolog (*rpmGC*). In the strains used in these studies, the L33* pseudogene has had the frameshift corrected (*rpmGC*^+^) so it encodes a functional, Zur-regulated L33* protein.

### Total cellular and ribosomal Zn concentration measurement by ICP-MS

Cells were grown in LB medium or MM to an OD_600_ of ~0.4, 5 ml and 500 ml cells from the same culture were harvested for measuring total cellular Zn content and ribosome associated Zn respectively. Cell numbers of the culture were quantified and crude ribosomes were purified as describe above. To measure total cellular Zn, four milliliter samples were collected before shock and at different time points after shock. Cells were washed twice with phosphate buffered saline (PBS) buffer containing 0.1 M EDTA followed by two chelex-treated PBS buffer only washes. Cells were then resuspended in 400 μl of chelex-treated PBS buffer from which 50 μl was used for OD_600_ measurement. 10 μl of 10 mg/ml lysozyme (dissolved in PBS) was added to the remaining cells and incubated at 37°C for 20 min. 600 μl of 5% HNO_3_ with 0.1% (v/v) Triton X-100 was added to the supernatant for total cellular samples or crude ribosome samples, which was boiled at 95°C for 30 min. After centrifuging the samples again, the supernatant was diluted with 1% HNO_3_. Zn levels were measured by ICP-MS (Perkin Elmer ELAN DRC II using ammonia as the reaction gas and gallium as an internal standard) and normalized against total cell numbers. Data represent mean ± SE of at least three separate experiments.

### Data availability

All data supporting the findings of this study are presented in the figures or available from the corresponding author upon reasonable request.

## Acknowledgements

We thank Dr. Ahmed Gaballa for providing the ZagA protein used in these studies. This work was supported by a grant from the National Institutes of Health (R35GM122461) to JDH.

## AUTHOR CONTRIBUTION

Conception, PC, XH and JDH; Designed and performed experiments, PC and XH; Manuscript drafted and edited, PC and JDH.

---

**Table S1:** Strains used in this study

| Strain | Genotype |
| --- | --- |
| **B. subtilis** |  |
| CU1065 | *trpC2 attSPβ sfp<sup>0</sup> [DHB(G)<sup>+</sup>]*, Lab stock |
| HB20101 | CU1065 Δ*folE2* |
| HB20102 | CU1065 Δ*purH* |
| HB20103 | CU1065 Δ*purB* |
| HB20104 | CU1065 Δ*purB* Δ*purH* |
| HB20105 | CU1065 Δ*znuABC* |
| HB20106 | CU1065 Δ*znuABC* Δ*purH* |
| HB20107 | CU1065 Δ*znuABC* Δ*purB* |
| HB20108 | CU1065 Δ*znuABC* Δ*purB* Δ*purH* |
| HB20117 | CU1065 *amyE*::*pfl* riboswitch-*lacZ* |
| HB20118 | CU1065 *amyE*::*pfl* riboswitch-*lacZ* Δ*folE2* |
| HB20119 | CU1065 *amyE*::*pfl* riboswitch-*lacZ* Δ*folE2* *thrC*::Pspac(hy)-*folE2* |
| HB20135 | CU1065 Δ*zagA* |
| HB20136 | CU1065 Δ*zagA* Δ*purH* |
| HB20137 | CU1065 Δ*zagA* Δ*purB* |
| HB20138 | CU1065 Δ*zagA* *amyE*::P<sub>spac</sub>(hy)-*zagA*(Bsu) |
| HB20139 | CU1065 Δ*zagA* *amyE*::P<sub>spac</sub>(hy)-*zagA*(Abay) |
| HB20140 | CU1065 Δ*zagA* *amyE*::P<sub>spac</sub>(hy)-*zagA*(Abau) |
| HB20141 | CU1065 Δ*rpmE* |
| HB20142 | CU1065 Δ*rpmEB* |
| HB20143 | CU1065 Δ*zagA* Δ*rpmE* |
| HB20144 | CU1065 Δ*zagA* Δ*rpmEB* |
| HB19657 | CU1065 *rpmGA*::*tet* *rpmGB*::*mls* *rpmE*::*spc* *rpmGC*<sup>+</sup> |
| HB19670 | CU1065 *rpmEB*::*tet* *rpmGC*<sup>+</sup> *sacA*::P<sub>folE2</sub>-*lux* |
| HB19672 | CU1065 *rpmE*::*spc* *rpmGC*<sup>+</sup> *sacA*::P<sub>folE2</sub>-*lux* |
| **E. coli** |  |
| DHP1 | DH1 (F<sup>-</sup>, *glnV44*(AS), *recA1*, *endA1*, *gyrA96* (Nal<sup>r</sup>), *thi1*, *hsdR17*, *spoT1*, *rfbD1*) |
| HE20101 | DHP1 pT18-empty, pT25 empty |
| HE20102 | DHP1 pT18 ZIP, pT25-ZIP |
| HE20103 | DHP1 pT18 *ZagA*(Bsu), pT25-FolE(Bsu) |
| HE20104 | DHP1 pT18 FolE(Bsu), pT25-ZagA(Bsu) |
| HE20105 | DHP1 pT18 *ZagA*(Abay), pT25 FolE(Abay) |
| HE20106 | DHP1 pT18 *ZagA*(Bsu), pT25 FolE(Abay) |
| HE20107 | DHP1 pT18 *ZigA*(Abay), pT25 FolE(Bsu) |
| HE20108 | DHP1 pT18 *ZigA*(Abau), pT25FolE(Abau) |

**Figure S1.**
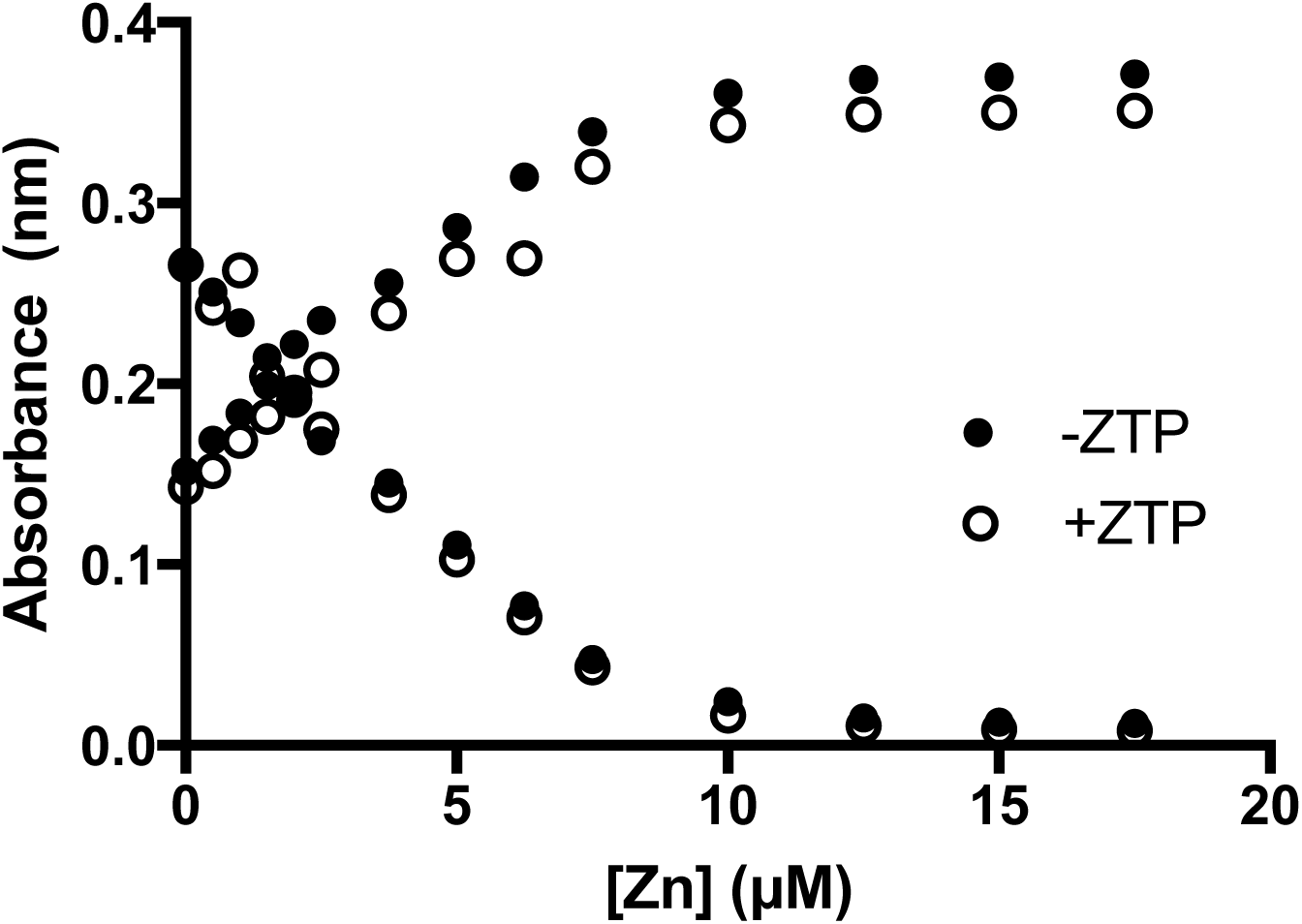
ZTP does not bind zinc with high affinity. Zinc was titrated into a mixture of 2 μM Magfura-2 without (filled circles) and with 10 μM ZTP (open circles). Absorbances at 325 nm (increasing values) and 366 nm (decreasing values) were plotted.

**Figure S2.**
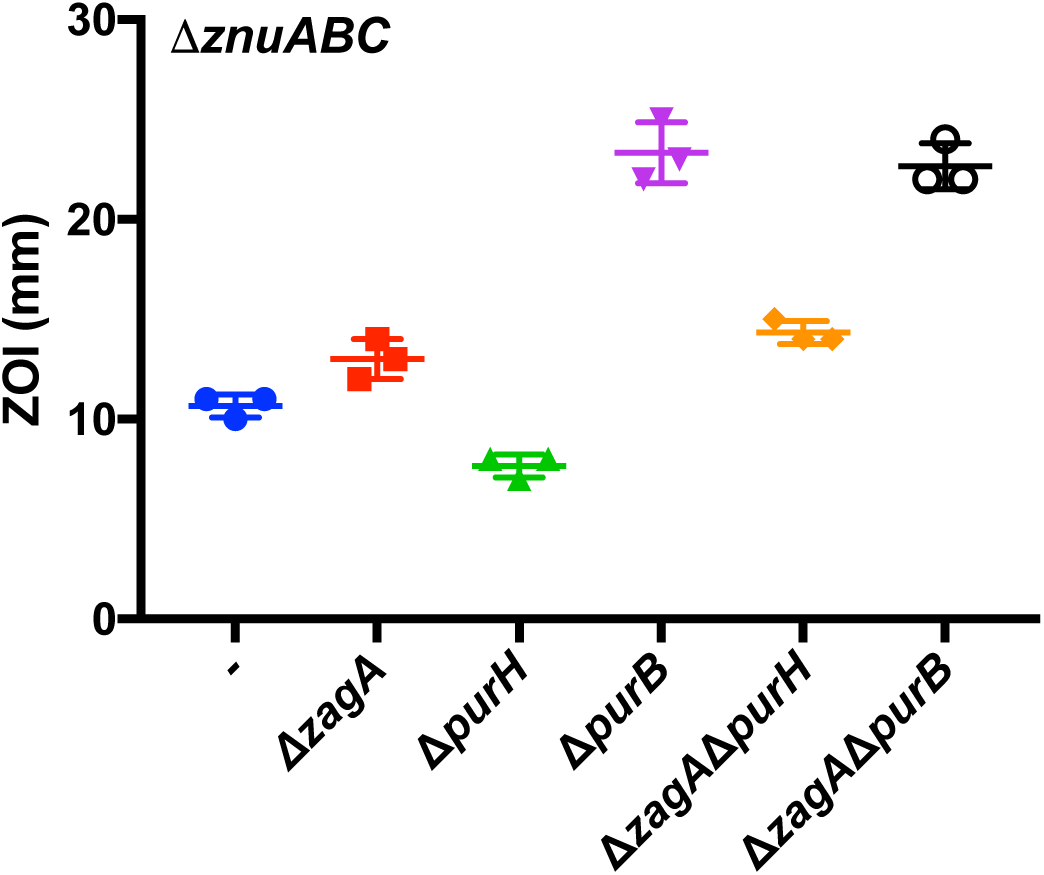
The protective effect of ZTP accumulation requires ZagA. EDTA sensitivity of *zagA*, *purH*, *purB* and *zagA purH*, and *zagA purB* mutants in a *znuABC* mutant background as measured by disk diffusion assay.

**Figure S3.**
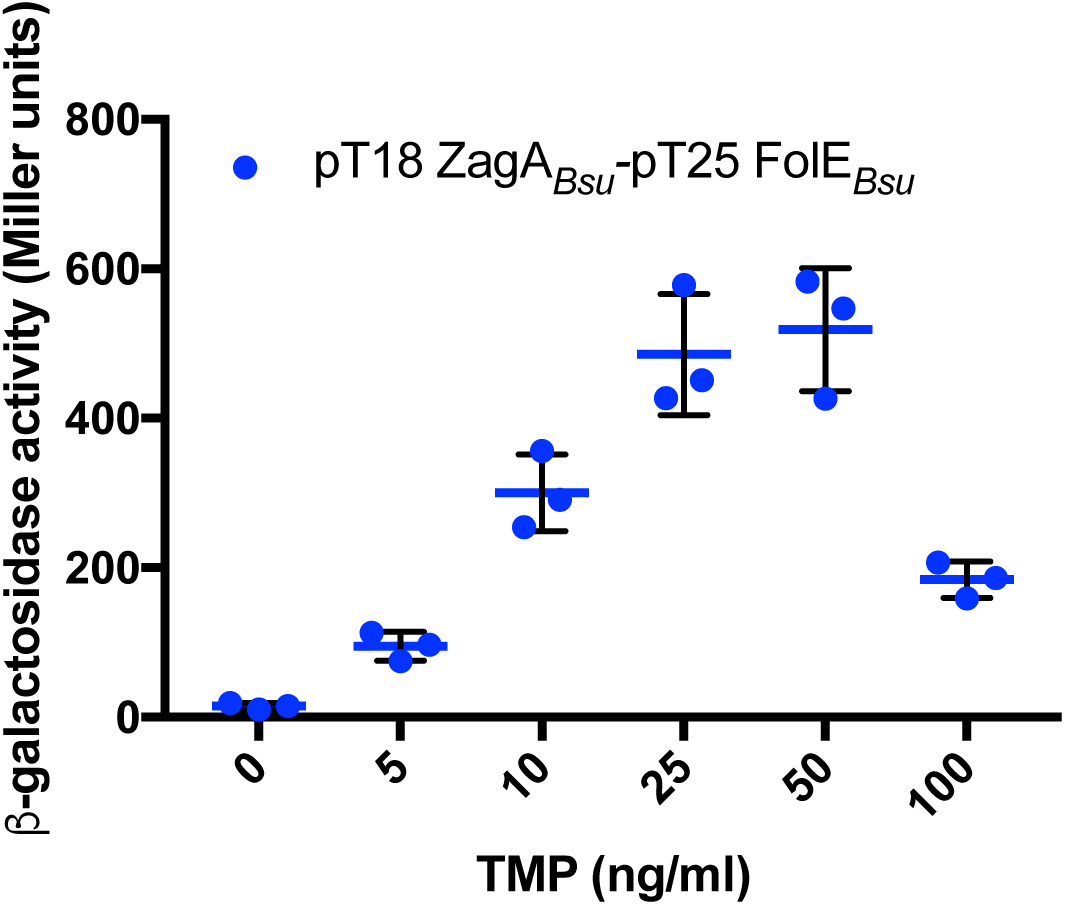
The ZagA-FolE interaction in response to trimethoprim is concentration dependent. *β*-galactosidase activity of the ZagA and FolE bacterial two hybrid constructs after 30 min of treatment with various concentrations of trimethoprim (TMP).

**Figure S4.**
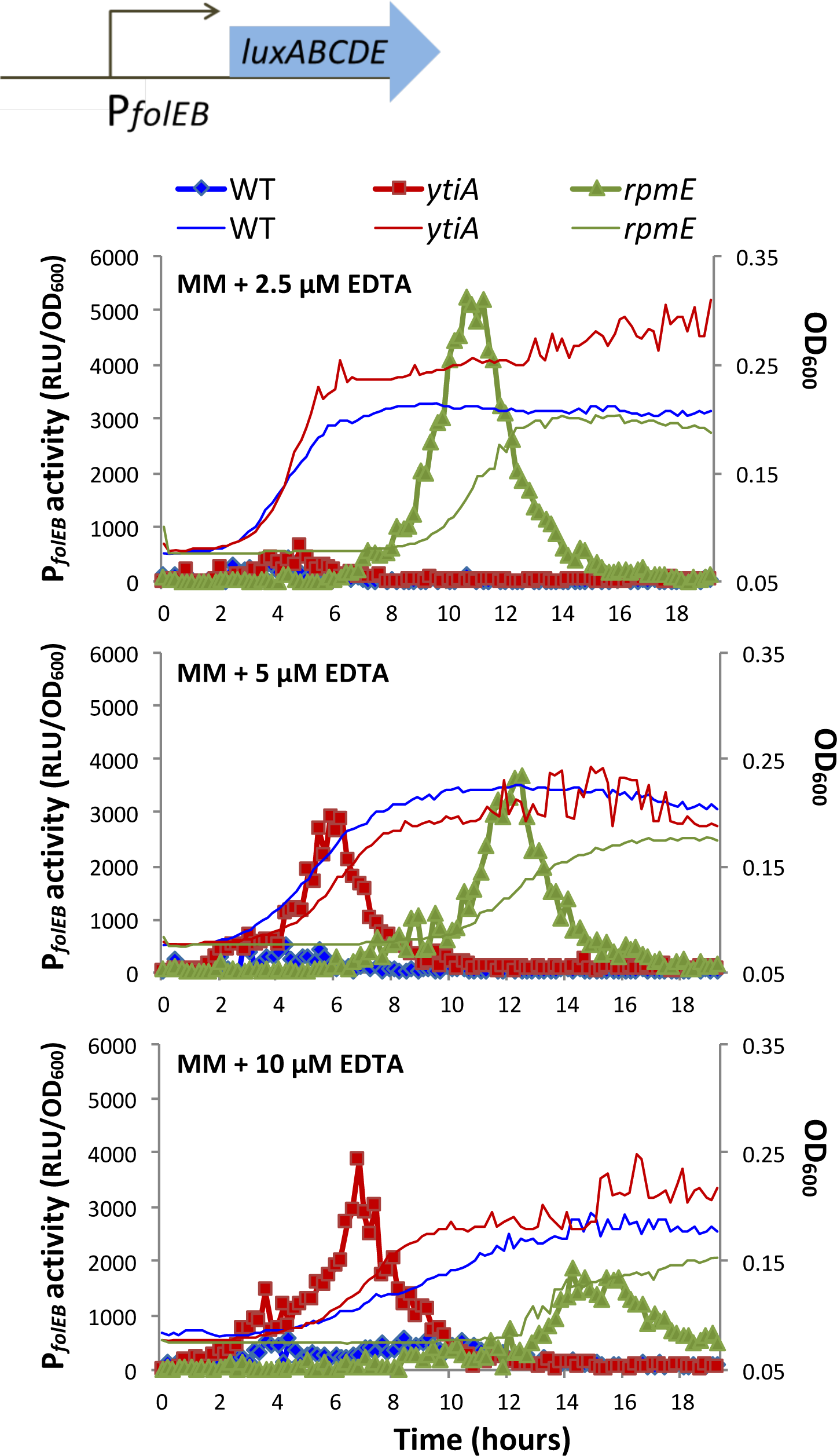
Zur regulon induction in response to zinc starvation is delayed in strains lacking the L31 (*rpmE*) or Zn-independent L31* (*ytiA*=*rpmEB*) ribosomal proteins. Growth curves of WT (blue), *rpmEB* (red) and *rpmE* (green) mutants in the presence of varying concentrations of EDTA. The induction of the *folEB* promoter-*lux* fusion in WT (blue diamonds), *rpmEB* (red squares), and *rpmE* mutants (green triangles) in the presence of varying concentrations of EDTA as a function of time is also shown.

